# Verification of nucleotide sequence reagent identities in original publications in high impact factor cancer research journals

**DOI:** 10.1101/2023.02.03.526922

**Authors:** Pranujan Pathmendra, Yasunori Park, Francisco J. Enguita, Jennifer A. Byrne

## Abstract

Human gene research studies that describe wrongly identified nucleotide sequence reagents have been mostly identified in journals of low to moderate impact factor, where unreliable findings could be considered to have limited influence on future research. This study examined whether papers describing wrongly identified nucleotide sequences are also published in high impact factor cancer research journals. We manually verified nucleotide sequence identities in original *Molecular Cancer* articles published in 2014, 2016, 2018 and 2020, including nucleotide sequence reagents that were claimed to target circRNAs. Using keywords identified in problematic 2018 and 2020 *Molecular Cancer* papers, we also verified nucleotide sequence identities in 2020 *Oncogene* papers that studied miRNA(s) and/or circRNA(s). Overall, 3.8% (253/6,647) and 4.3% (50/1,165) nucleotide sequences that were verified in *Molecular Cancer* and *Oncogene* papers, respectively, were found to be wrongly identified. These wrongly identified nucleotide sequences were distributed across 18% (92/500) original *Molecular Cancer* papers, including 38% *Molecular Cancer* papers from 2020, and 40% (21/52) selected *Oncogene* papers from 2020. Original papers with wrongly identified nucleotide sequences were therefore unexpectedly frequent in two high impact factor cancer research journals, highlighting the risks of employing journal impact factors or citations as proxies for research quality.

## Introduction

Despite technological advances, growing research workforce capacity and billion-dollar budgets devoted to biomedical research in first-world countries, biomedical research translation continues to fall short of the expectations generated by research investments [1]. Inefficient research translation is fueled by the reproducibility crisis, where many pre-clinical research results cannot be independently reproduced [2–4]. The emphasis upon the publication of positive findings has likely led to the publication of false-positive results [2, 5, 6]. Where these results are not reproduced by other studies, these contradictory or discordant results may be less likely to be reported, leading to a growing problem of falsely-positive research results in the biomedical literature [5, 6].

While most incorrect pre-clinical research is believed to derive from genuine research [7], some irreproducible research results may reflect data falsification and fabrication [8, 9]. The incidence of research fraud is likely to be underestimated, not least because research fraud is actively concealed in publications [8]. Over the past several years, the analysis of research fraud has shifted from focusing on research fraud perpetrated by individuals, to include research fraud that may be enabled by research contracting cheating organizations or paper mills [10–14]. Paper mills represent hidden organizations that allegedly supply a range of undeclared research services to potential authors [10, 13–16]. Undeclared paper mill services can include the provision of research datasets that paper mill clients can include in manuscripts, and complete manuscripts that can be authored by individuals or teams as required [10, 14, 17].

There is growing evidence that human genes may be targeted by paper mills for the production of preclinical research manuscripts [11, 15, 18]. Several studies have identified repetitive gene and cancer research publications that feature unusual levels of structural and textual similarity, including implausibly superficial, novelty-based research justifications, and the repeated application of generic experimental approaches [15, 18–21]. The mass retraction of human gene research papers in response to manipulated peer review [22] was linked with some authors disclosing the use of paper mill services [23]. More recently, numerous biomedical and chemistry journals have published editorials recognizing manuscript submissions from paper mills [24–38], where some editorials have recognized papers that examined human genes [28, 29, 31, 38]. Some journals have undertaken retractions of papers that are suspected to have originated from paper mills [28-30, 35, 36]. A rapid scoping review published in 2022 identified over 300 papers that had been retracted in response to paper mill involvement, where the keyword ‘miR” (microRNA) was the most frequently identified key word [39].

The rapid production of many gene research manuscripts at minimal cost could provide limited time for quality control, which could result in errors such as wrongly identified nucleotide sequence reagents [11, 18]. The semi-automated tool Seek & Blastn was created to verify the identities of published nucleotide sequence reagents that are claimed to target human genes and transcripts [19], where the application of Seek & Blastn has demonstrated the widespread occurrence of wrongly identified nucleotide sequence reagents in repetitive human gene research papers [19–21]. Our most recent application of Seek & Blastn screened over 11,700 original human research papers and identified 712 papers that described wrongly identified nucleotide sequence(s) [21]. Seek & Blastn screening of original papers in the journals *Gene* and *Oncology Reports* revealed that yearly proportions of original papers with wrongly identified sequence(s) ranged from 0.5-4.2% and 8.3-12.6%, respectively [21].

Most human gene research papers with features of paper mill support have been identified in journals of low to moderate impact factor (IF) [18–21]. This finding is likely to at least partly reflect the negatively skewed distribution of journal IF’s [40, 41]. For example, high IF cancer research journals have been defined as having an IF ≥ 7.0 [42], which corresponds to ∼20% of cancer research journals. While recognizing the limited utility of journal IF as a measure of research quality [41], the perceived significance of problematic human gene research papers could be discounted through their publication in lower IF journals.

Despite paper mills being considered a problem of lower IF journals, papers with features of paper mill involvement have also been published in high IF journals, as recognized by editorials in the journals *International Journal of Cancer*, *Molecular Therapy* and *Molecular Therapy-Nucleic Acids* [34–36]. Our team has also described human gene research papers with wrongly identified nucleotide sequences that were published in high IF journals [19, 21]. Nonetheless, it is unclear whether low numbers of problematic human gene research papers in high IF journals [19, 21] simply reflect low numbers of high IF journals [40, 41], and/or that few problematic human gene research papers have been published by high IF journals.

We have therefore undertaken a literature screening approach to examine the frequency of human gene research papers with wrongly identified nucleotide sequence reagents in two high IF cancer research journals, as judged by 2019 journal IF (https://clarivate.com/) [42]. We chose to examine *Molecular Cancer*, an online, open-access journal published by BMC (Springer Nature), as Seek & Blastn screening of keyword-driven literature corpora had identified problematic *Molecular Cancer* papers published in 2014 [21]. Although *Molecular Cancer* was not a high IF journal in 2014 (IF=4.3), *Molecular Cancer* has experienced a marked rise in journal IF, reaching IFs of 15.3 in 2019, 27.4 in 2020, and 41.4 in 2021 (Fig. 1). As a result, *Molecular Cancer* was the 3rd-ranked molecular biology and biochemistry journal in 2020 and 2021, following only *Nature Medicine* and *Cell*. We also verified nucleotide sequence reagent identities in a selected corpus of 2020 *Oncogene* papers. *Oncogene* is published by Springer Nature under a hybrid open-access/ subscription publication model. Unlike *Molecular Cancer*, *Oncogene* has shown a relatively stable journal IF ranging from 6.6-9.9 during 2014-2021 (Fig. 1).

**Figure 1.**
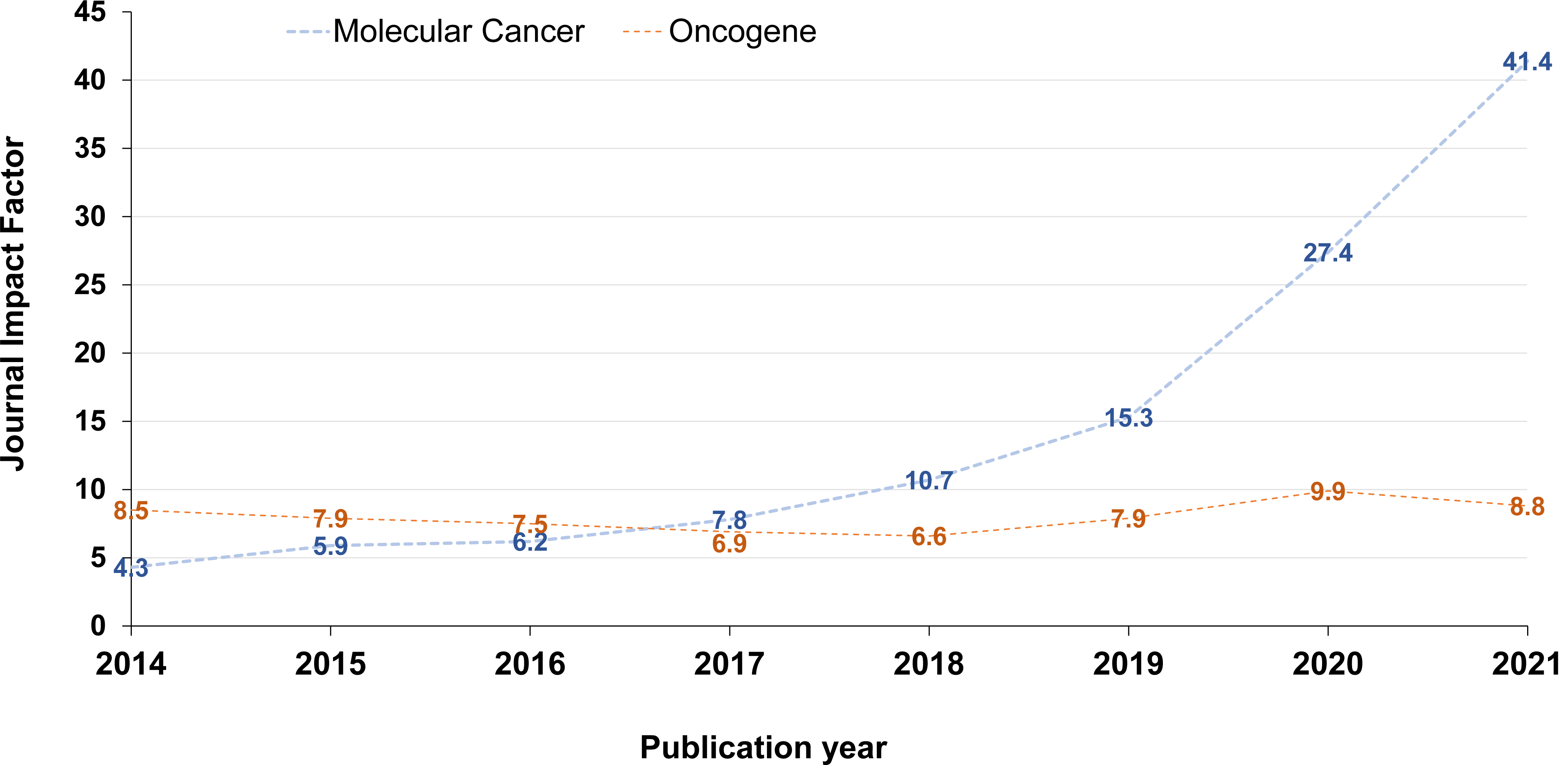
Journal impact factors (https://clarivate.com/) (Y axis) for *Molecular Cancer* (blue) and *Oncogene* (orange) from 2014-2021 (X axis). Journal impact factors have been rounded to one decimal place.

As most *Molecular Cancer* papers described nucleotide sequence reagents in supplementary files and not in the publication text, these papers proved to be unsuitable for Seek & Blastn screening [19]. We therefore manually verified the identities of all nucleotide sequence reagents that were claimed to target unmodified human gene targets in original *Molecular Cancer* papers published in 2014, 2016, 2018 and 2020. These publication years were chosen so that proportions of problematic *Molecular Cancer* papers could be compared with those identified in *Gene* and *Oncology Reports* in 2014, 2016 and 2018 [21]. As some *Molecular Cancer* papers described nucleotide sequence reagents that were claimed to target human circular RNA (circRNA) transcripts, we developed protocols to verify the identities of circRNA-targeting reagents. Using keywords identified in some problematic *Molecular Cancer* papers (miRNA, miR, circular RNA, or circRNA), we undertook keyword-driven searches of all original 2020 *Oncogene* papers. We manually verified the identities of all nucleotide sequence reagents that were claimed to target unmodified human gene targets in all 2020 *Oncogene* papers that referred to microRNAs and/or circRNAs. As we will describe, these analyses identified unexpectedly high proportions of human gene research papers with wrongly identified nucleotide sequences in two high IF cancer research journals. Our results therefore indicate that problematic human gene research publications may be unexpectedly frequent in some high IF cancer research journals.

## Results

### Molecular Cancer corpus

In total, 500 original *Molecular Cancer* papers were published in 2014, 2016, 2018 and 2020 (Table 1), where numbers of original papers ranged from 59 papers in 2016, to 249 papers in 2014 (Fig. 2A). Most (334/500, 67%) original *Molecular Cancer* papers were included for analysis as they described human research and included at least one nucleotide sequence that was claimed to target a non-modified human gene or genomic sequence (Fig. 2A, Table 1). The proportions of *Molecular Cancer* papers that met the study inclusion criteria ranged from 29/59 (49%) in 2016 to 74/82 (90%) in 2020 (Fig. 2A).

**Table 1.** Molecular Cancer and Oncogene corpora that were screened for wrongly identified nucleotide sequence reagents

|  | *Molecular Cancer* | *Oncogene* |
| --- | --- | --- |
| Corpus description | All original papers published in 2014, 2016, 2018 and 2020 | Keyword driven search of original papers published in 2020 |
| Number of original papers screened | 500 | 52 |
| Proportion (percentage) of all original papers that were eligible for analysis | 334/500 (67%) | 42/52 (81%) |
| Total number of nucleotide sequence reagents eligible for fact-checking | 6,647 | 1,165 |
| Number of nucleotide sequence reagents per article, median (range) | 13 (1-153) | 20 (2-115) |
| Proportion (percentage) of wrongly identified nucleotide sequence reagents | 253/6,647 (3.8%) | 50/1,165 (4.3%) |
| Percentage (proportion) of problematic papers | 92/500 (18%) | 21/52 (40%) |
| Number of wrongly identified nucleotide sequences per article, median (range) | 2 (1-14) | 2 (1-5) |
| Proportion (percentage) of wrongly identified nucleotide sequence reagents according to error types: | 253/253 (100%) | 50/50 (100%) |
| Claimed targeting reagents that were predicted to target a different human gene or genomic sequences | 137/253 (54%) | 27/50 (54%) |
| Claimed targeting reagents that were predicted to be non-targeting in human | 114/253 (45%) | 23/50 (46%) |
| Claimed non-targeting reagents that were predicted to target a human gene | 2/253 (0.8%) | 0/50 (0%) |

**Figure 2.**
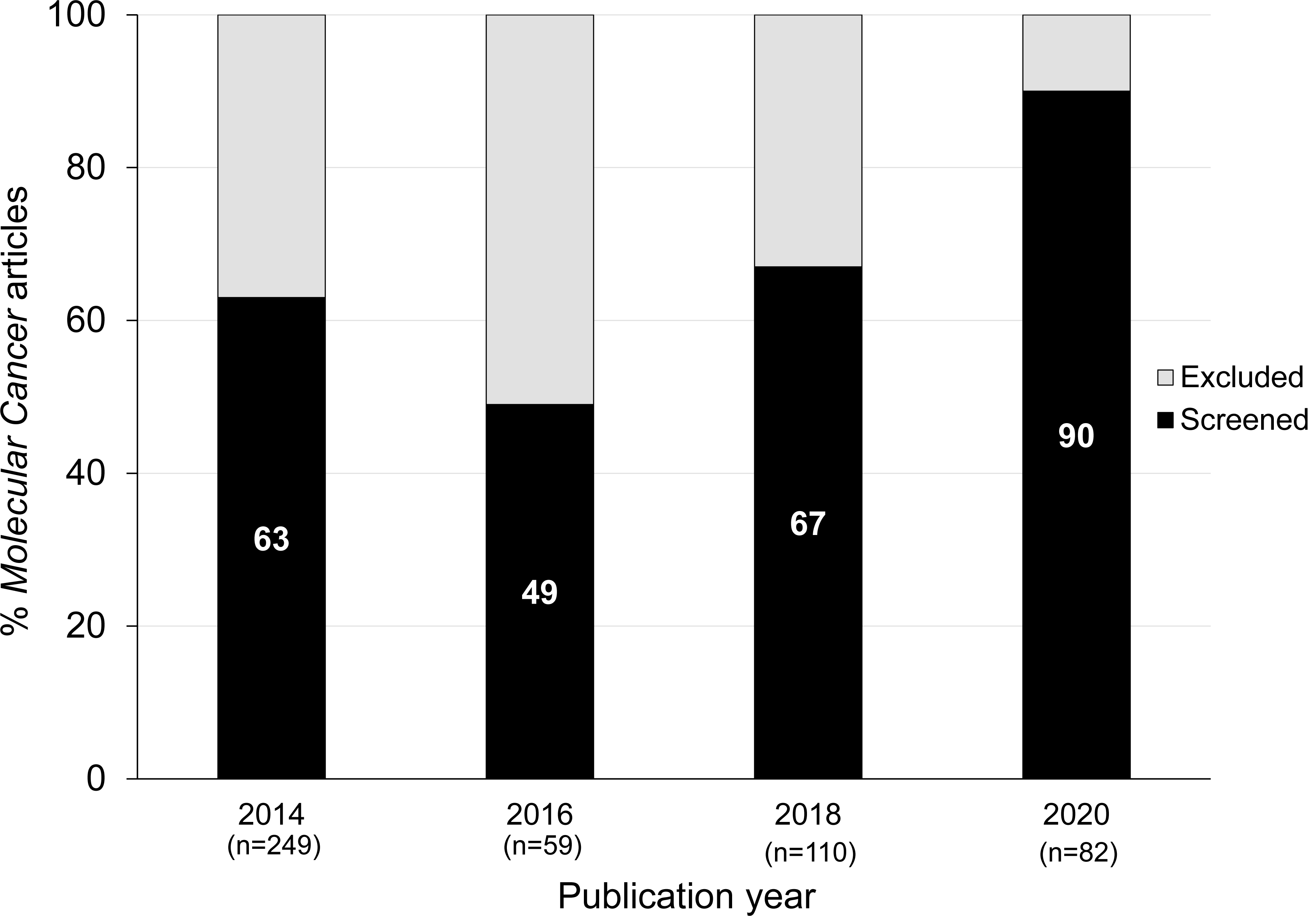
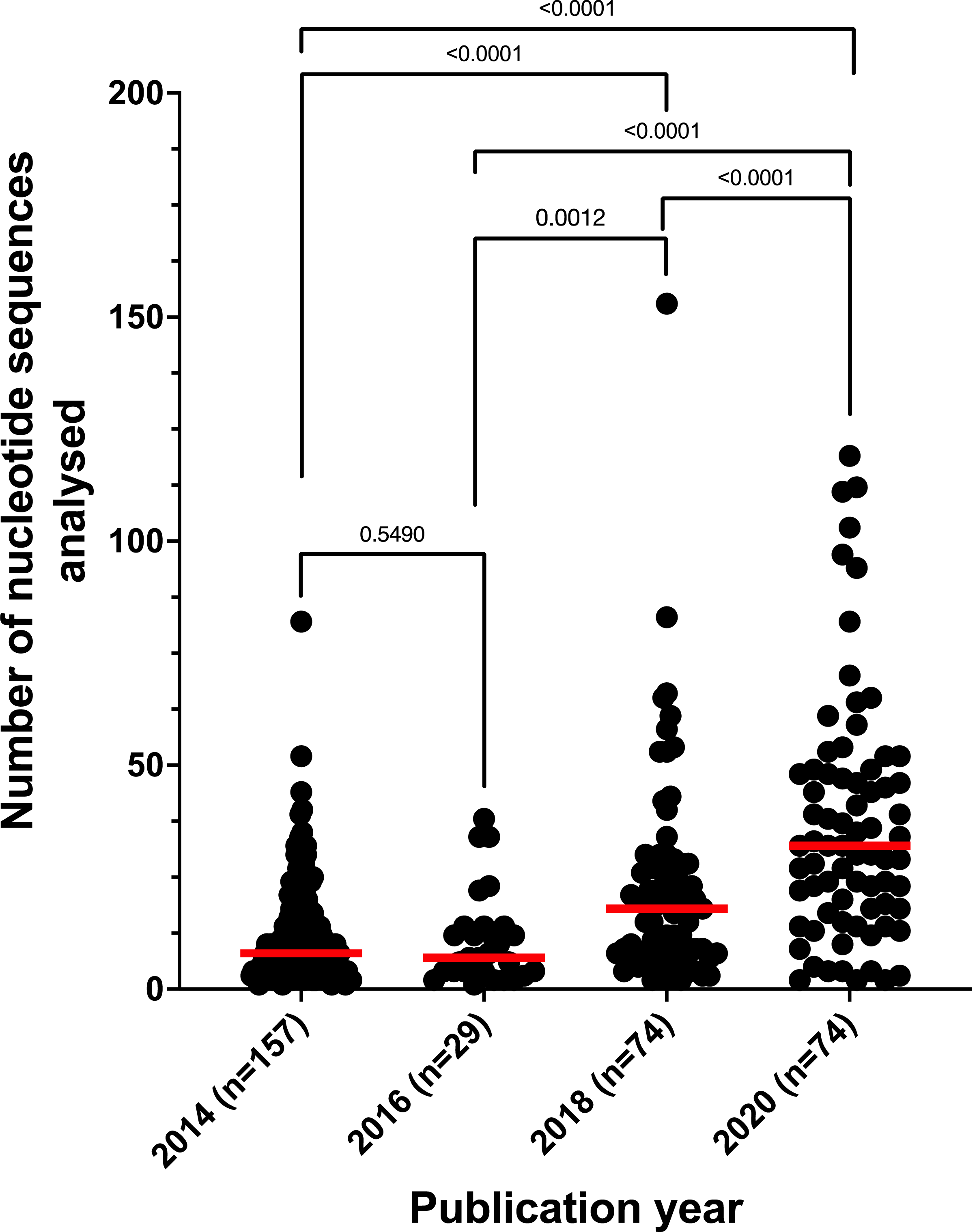
Summary of original papers published in *Molecular Cancer* in 2014, 2016, 2018 and 2020. Numbers of original *Molecular Cancer* papers (analysed) per year are shown below the X-axis. **A**) Percentages of original *Molecular Cancer* papers (Y-axis) that were either screened (black, percentage values shown in white text) or excluded from analysis (gray) per year (X-axis). **B**) Numbers of nucleotide sequences per *Molecular Cancer* paper (Y-axis) according to publication year (X-axis). Only original *Molecular Cancer* papers that described at least one nucleotide sequence reagent were included in these analyses. Individual/ median numbers of nucleotide sequences/ paper are shown as black dots/ red horizontal lines, respectively. The Mann-Whitney test was employed to compare median nucleotide sequence numbers/ paper according to publication year, as indicated by p values.

The 334 *Molecular Cancer* papers included 6,647 nucleotide sequences, with a median of 13 nucleotide sequences/ paper (range 1-153) (Table 1). The numbers of nucleotide sequence reagents per paper progressively increased from 2014-2020 (Fig. 2B). For example, the median number of nucleotide sequences per paper increased from 8 sequences/ paper in 2014, to 32 sequences/ paper in 2020 (Mann-Whitney test, p<0.0001, n=231) (Fig. 2B).

## Problematic *Molecular Cancer* papers describing wrongly identified nucleotide sequence(s)

Whereas no 2014 or 2016 *Molecular Cancer* papers described nucleotide sequences that were claimed to target human circular RNAs (circRNAs), 39 *Molecular Cancer* papers in 2018 and 2020 described circRNA-targeting reagents. As we had not previously verified the identities of circRNA-targeting reagents, new protocols were developed to recognize the particular targeting requirements of some circRNA reagents (Figs. 3, 4, see Methods).

**Figure 3.**
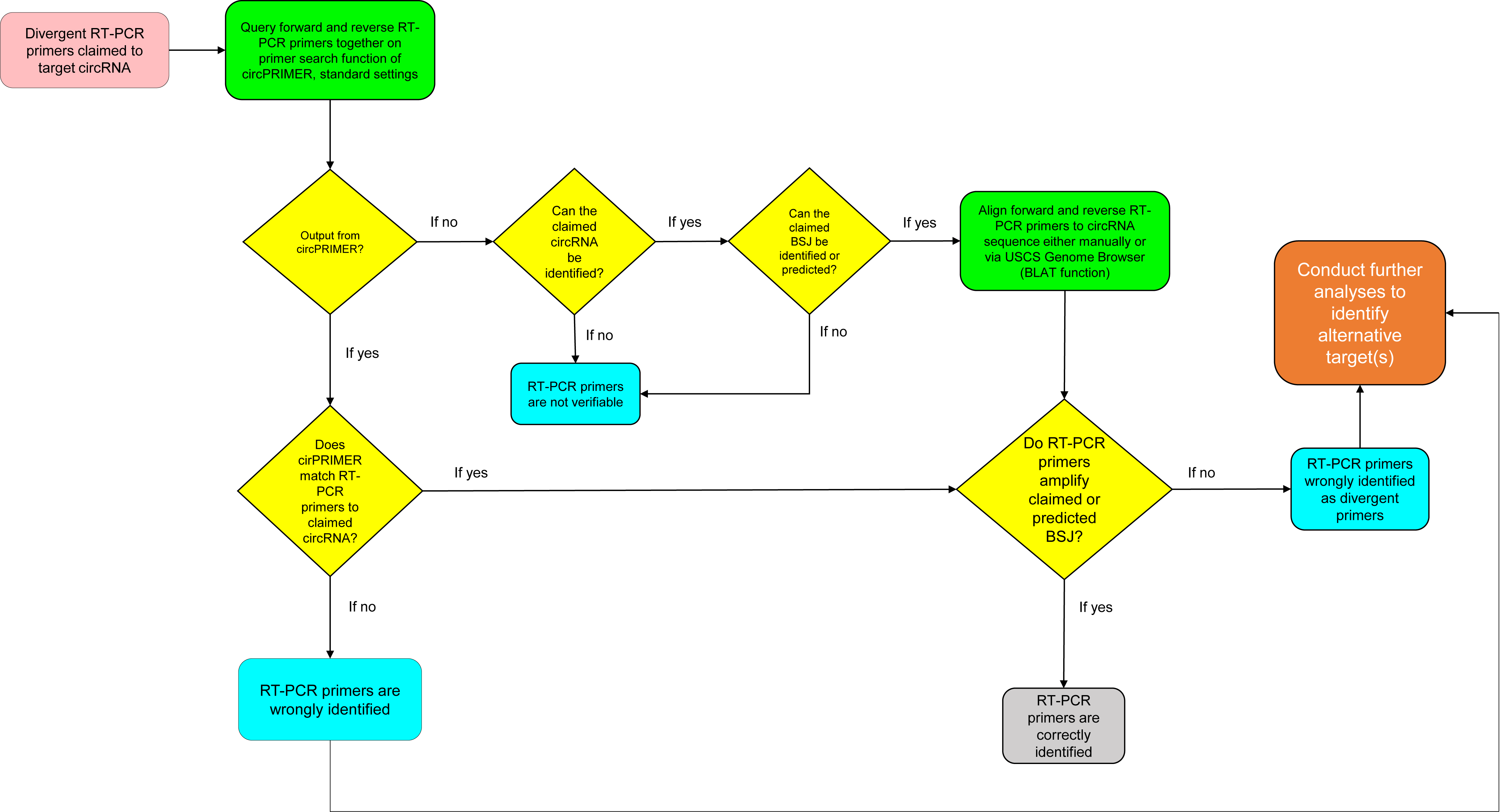
Flow chart summarising the workflow that was used to manually verify the identities of divergent RT-PCR primers claimed to target human circRNAs.

**Figure 4.**
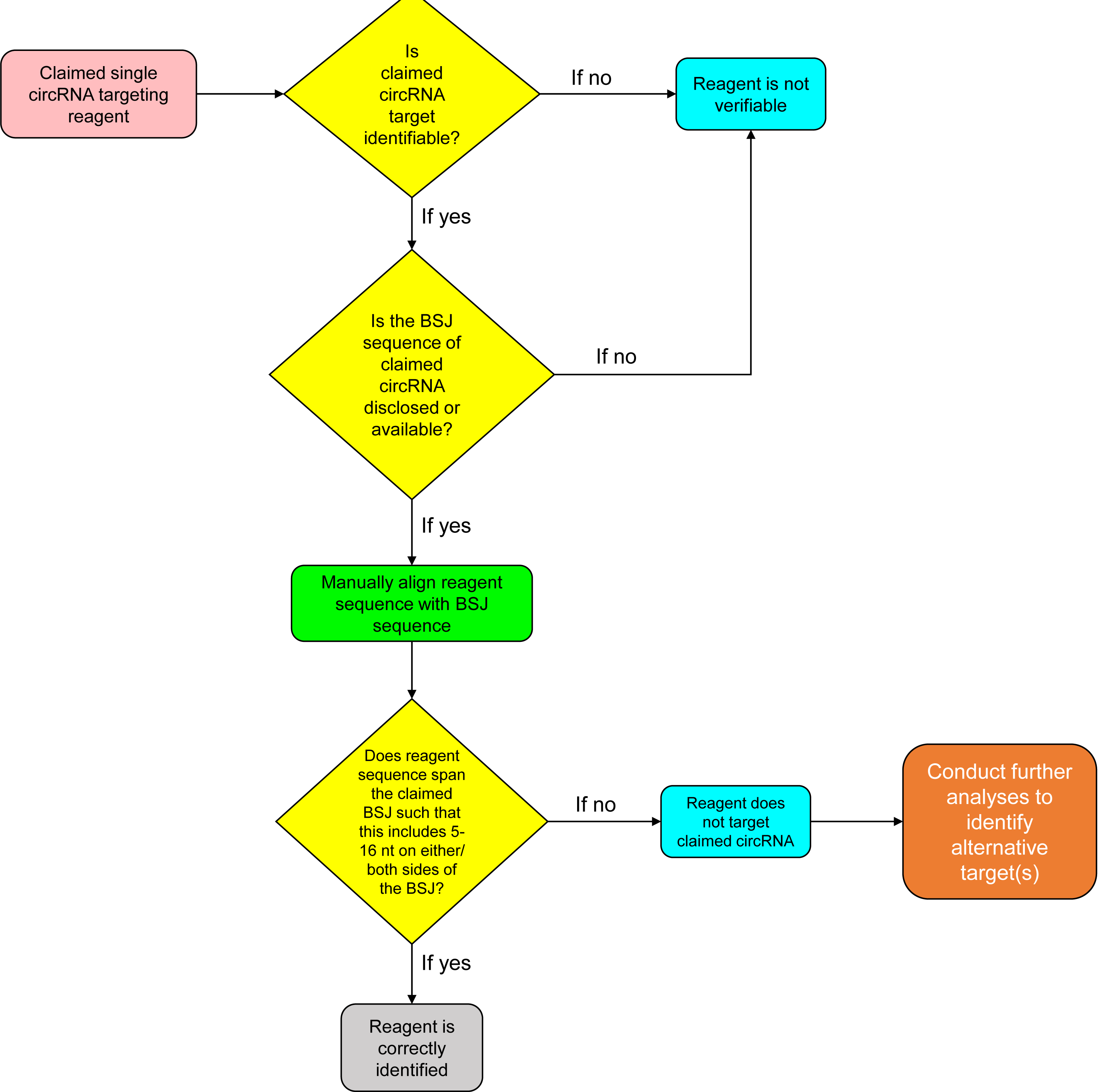
Flow chart summarising the workflow that was used to manually verify the identities of single nucleotide sequence reagents (siRNAs, shRNAs, other oligonucleotide probes) claimed to target human circRNAs.

Of the 6,647 nucleotide sequences whose identities were manually verified, 253 (3.8%) nucleotide sequences were predicted to be wrongly identified (Table 1, Fig. 5A, Table S1). Similar proportions of incorrect sequences represented targeting reagents that were either verified to target a different human gene or genomic sequence (137/253, 54%), or predicted to be non-targeting in human (114/253, 45%) (Table 1, Fig. 5B). In contrast, very few (2/253, 0.8%) wrongly identified sequences represented claimed non-targeting si/shRNA reagents that were instead predicted to target a human gene (Table 1, Fig. 5B).

**Figure 5.**
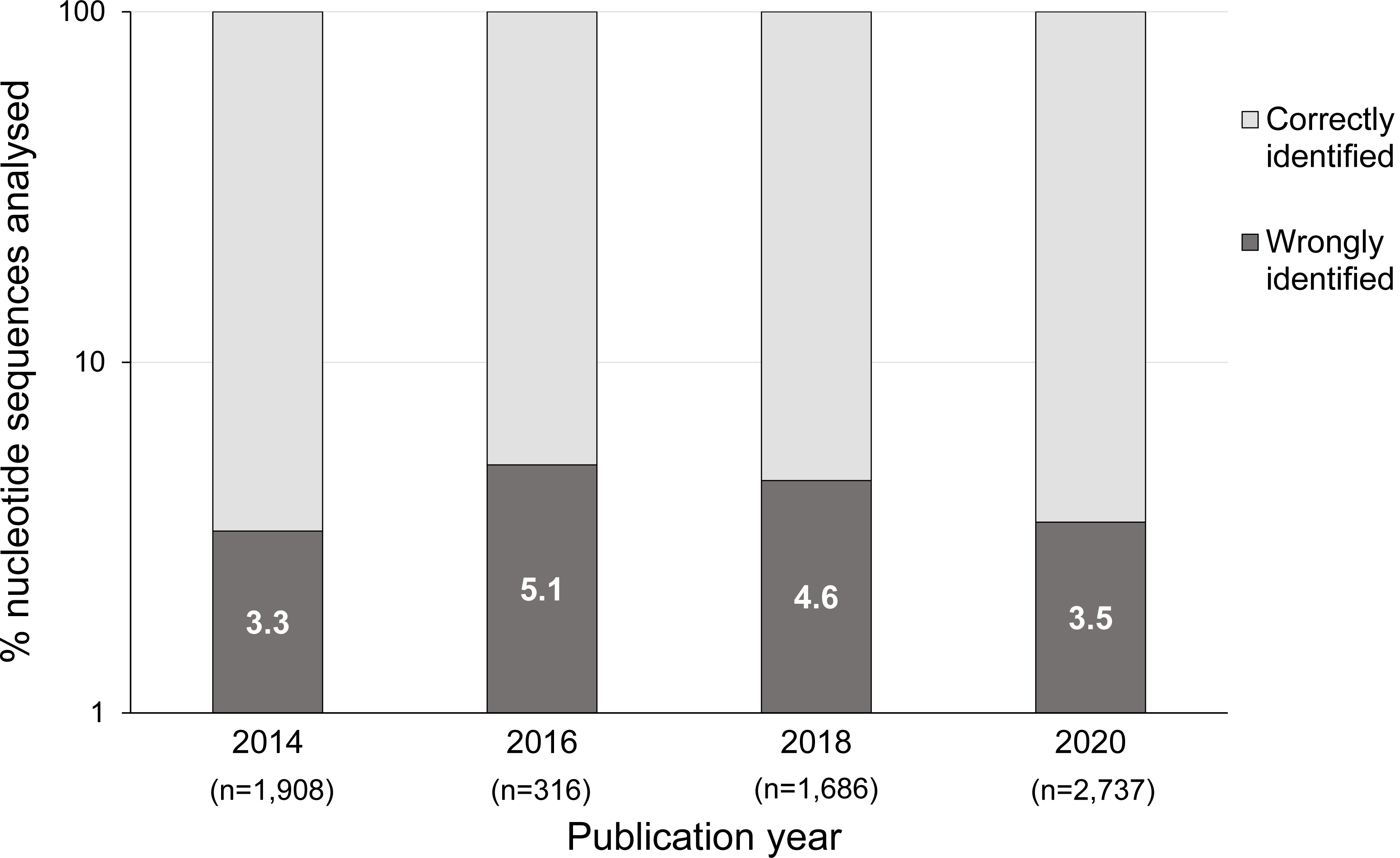
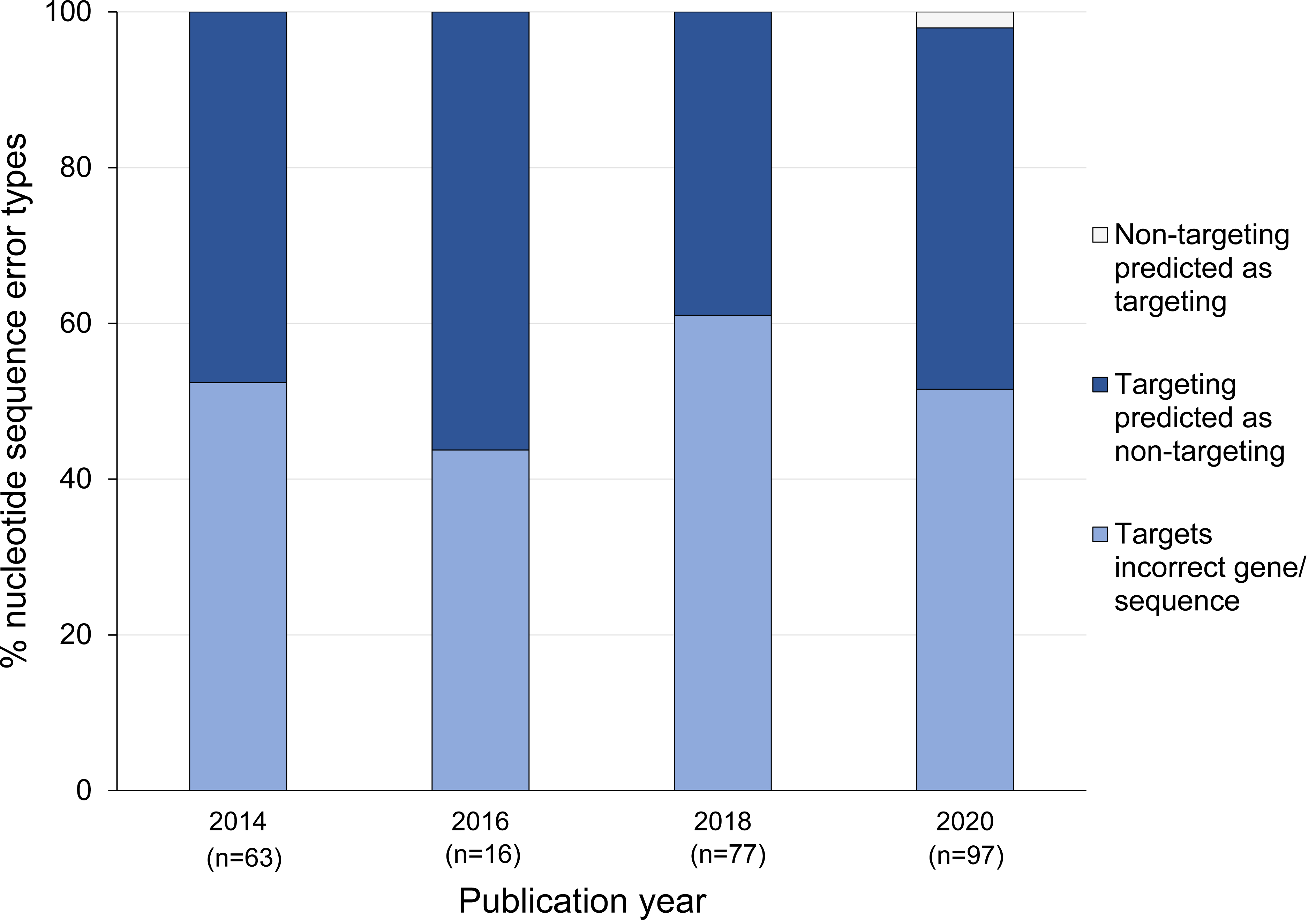
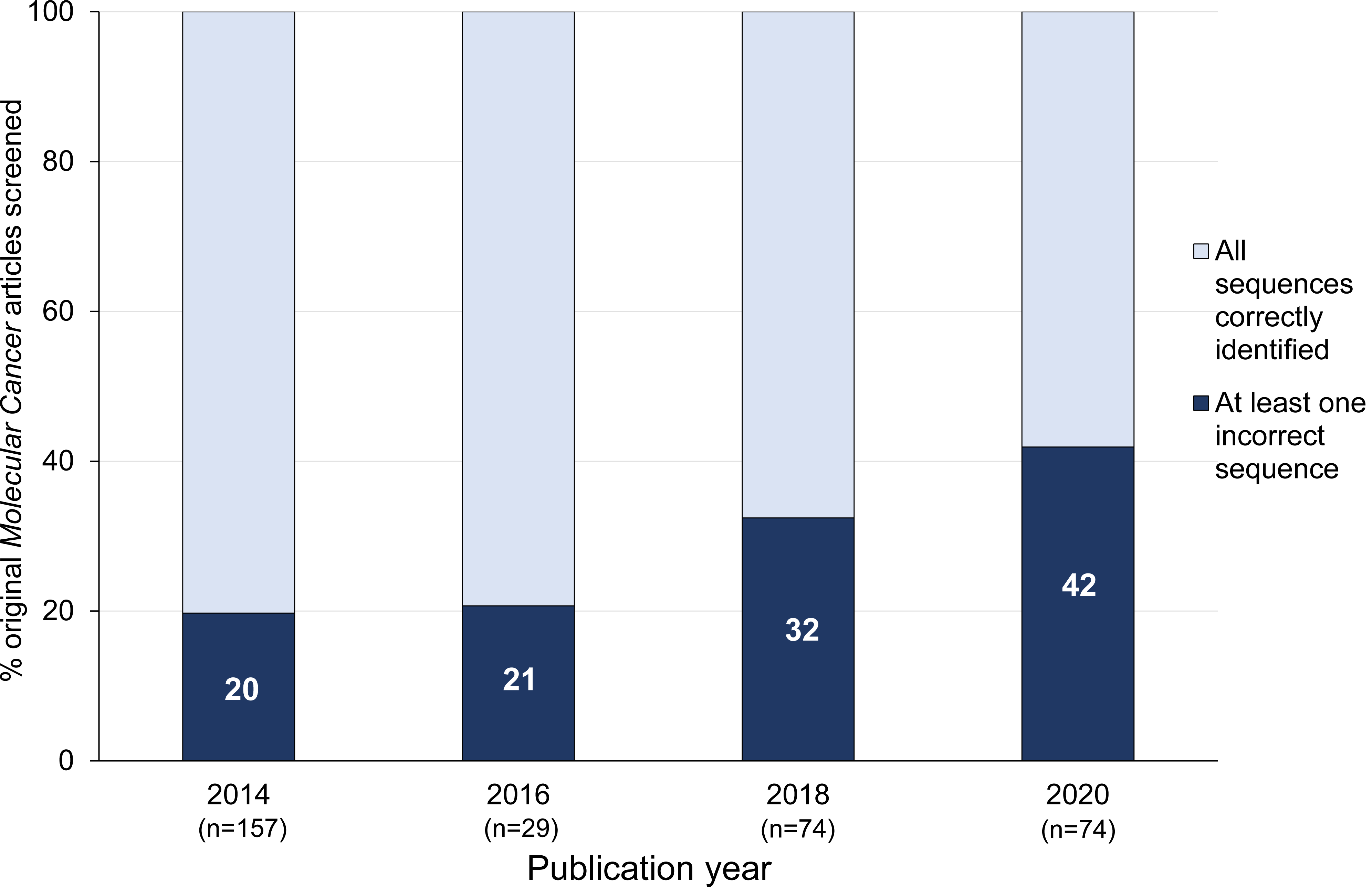
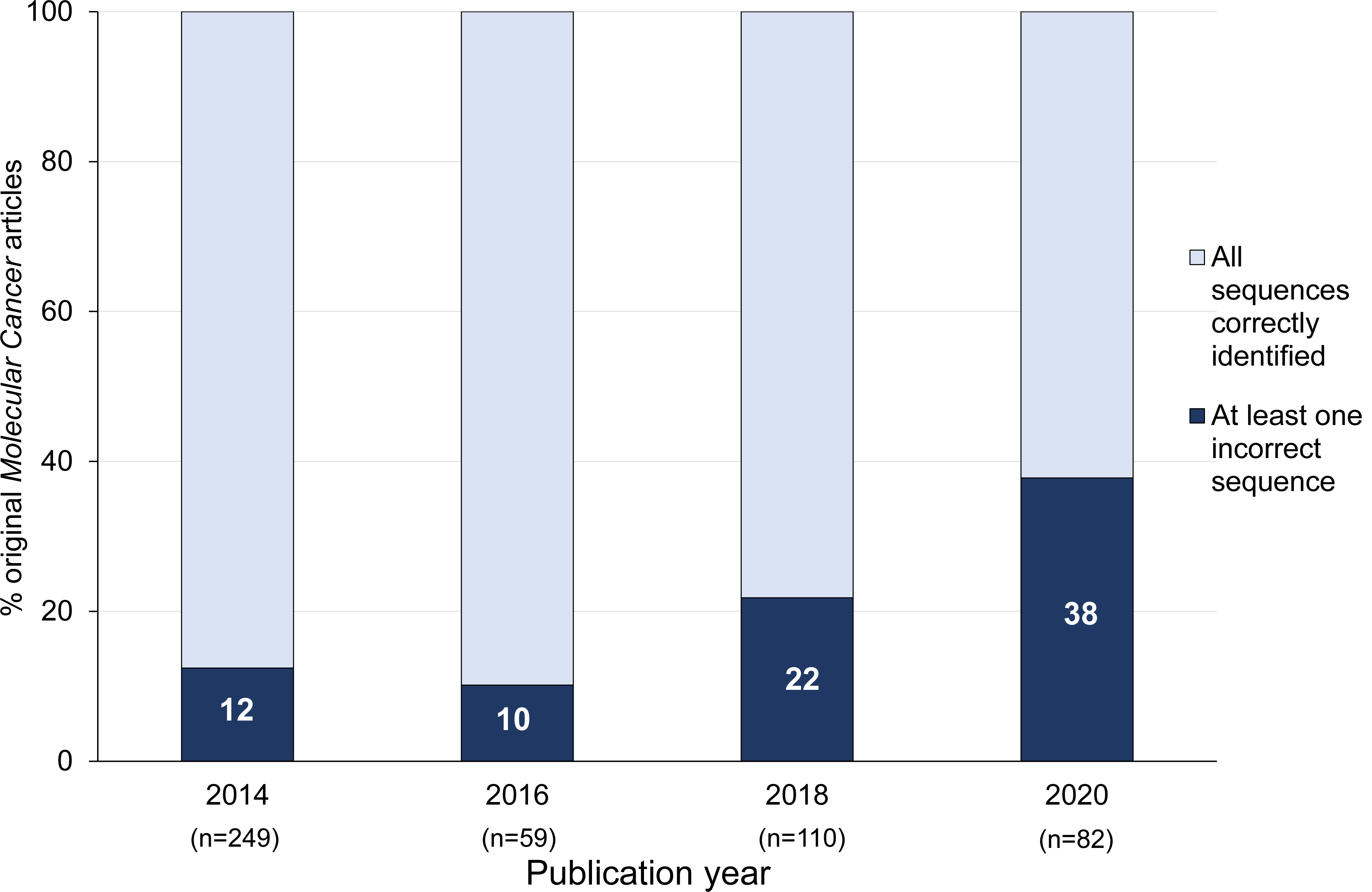
Summary of original *Molecular Cancer* papers in 2014, 2016, 2018 and 2020 that described at least one wrongly identified nucleotide sequence. **A**) Percentages of nucleotide sequences (Y axis, log scale) that were correctly (light gray) or wrongly identified (dark grey, percentages shown in white text) per publication year (X axis). Numbers of nucleotide sequences analysed in *Molecular Cancer* papers per year are shown below the X axis. **B**) Percentages of wrongly identified nucleotide sequences according to nucleotide sequence identity error types (Y axis) and publication year (X axis). Nucleotide sequence identity error types are shown as follows: claimed targeting reagents predicted to target a different gene or sequence (mid blue); claimed targeting reagents predicted to be non-targeting in human (dark blue); claimed non-targeting reagents predicted to target a human gene (light gray). Numbers of wrongly identified nucleotide sequences per publication year are shown below the X-axis. **C**, **D**) Percentages of screened (**C**) or original *Molecular Cancer* papers (**D**) (Y axes) that described at least one wrongly identified reagent (dark blue, percentages shown in white text) versus all other papers (light blue), according to publication year (X axis). Numbers of papers per year are shown below the X axis.

The 253 wrongly identified nucleotide sequences were distributed across 92/334 (28%) screened *Molecular Cancer* papers (Fig. 5C) and 92/500 (18%) original *Molecular Cancer* papers (Table 1, Fig. 5D, Table S2). These 92 papers included 3 *Molecular Cancer* papers from 2014 that were reported to describe wrongly identified nucleotide sequence(s) [19, 21]. Proportions of problematic papers ranged from 6/59 (10%) in 2016 to 31/82 (38%) in 2020 (Fig. 5D). The median number of wrongly identified sequences/paper was 2 (range 1-14) (Table 1, Fig. 6), where the numbers of wrongly identified and analysed sequences per paper were significantly positively correlated (Spearman’s rho=0.2085, 95% Cl=-0.0023-0.4016, p=0.0461, n=92).

**Figure 6.**
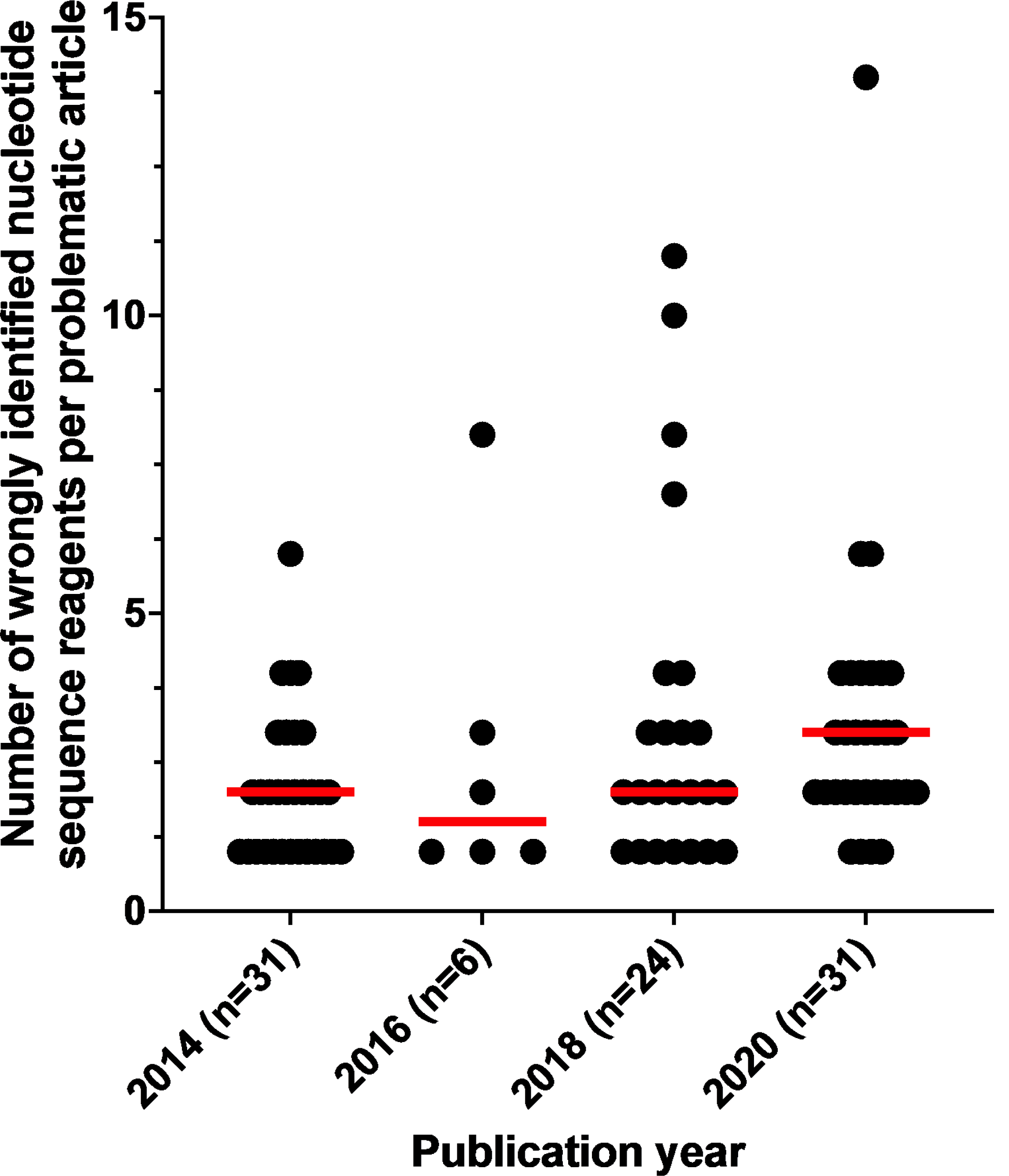
Numbers of wrongly identified nucleotide sequence reagents in problematic *Molecular Cancer* papers (Y axis) according to publication year (X axis). Individual/ median numbers of wrongly identified nucleotide sequences/ paper are shown as black dots/ red horizontal lines, respectively. Numbers of problematic *Molecular Cancer* papers per publication year are shown below the X axis.

The 92 problematic *Molecular Cancer* papers described experiments in human cancer models corresponding to 26 cancer types, most frequently gastric, colorectal or non-small cell lung cancer (Table S2). Almost all (85/92, 92%) problematic papers analysed a single cancer type. Problematic papers described a median of 2 genes or transcripts in their titles (range 0-7) (Table S2). Most problematic paper titles (79/92, 86%) mentioned at least one protein-coding gene. Just over half of titles (49/92, 53%) mentioned non-coding RNA(s) (ncRNAs), which were typically miR(s) (32/49, 65%) or circRNA(s) (16/49, 33%). Whereas most 2014 titles mentioned only protein-coding gene(s) (22/31, 71%), most 2020 titles combined protein-coding gene(s) and ncRNA(s) (22/31, 71%), which were again typically miR(s) (12/22, 55%). Sixteen problematic papers that referred to circRNAs in their titles were published in 2018 and 2020, where titles typically combined circRNAs with protein-coding gene(s) and/or miR(s) (14/16, 88%) (Table S2).

### Wrongly identified or non-verifiable reagents for the analysis of human circRNAs

Eleven problematic *Molecular Cancer* papers described 23 wrongly identified reagents that were claimed to target circRNAs (Table 2, Table S1). These claimed circRNA-targeting reagents were predicted to either target different human transcripts from those claimed (17/23, 74%) or to be non-targeting in human (6/23, 26%) (Table 2). Wrongly-identified circRNA-targeting sequences included claimed divergent RT-PCR primers that were predicted to amplify linear transcripts, and single reagents that showed significant identity to linear transcripts (see Methods, Table 2, Table S1). The identities of a further 27 circRNA-targeting reagents could not be verified (Table 2), either because the claimed circRNA sequence could not be identified in external databases, or in the case of single reagents, because the back splice junction (BSJ) sequence was not provided (see Methods, Tables S3-S5). Non-verifiable circRNA targeting reagents were identified in 4 problematic *Molecular Cancer* papers (Tables S3, S5). An additional 5 *Molecular Cancer* papers included non-verifiable circRNA targeting reagents, where all other nucleotide sequences appeared to be correctly identified (Tables S4, S5).

**Table 2.** Wrongly identified and non-verifiable nucleotide sequence reagents that were claimed to target human circRNAs in Molecular Cancer and Oncogene papers

| **Claimed circRNA targeting reagents** | ***Molecular Cancer*** | ***Oncogene*** |
| --- | --- | --- |
| **Wrongly identified** human circRNA targeting reagents | 23/23 (100%) | 6/6 (100%) |
| circRNA targeting reagents predicted to **target a different human transcript** | 17/23 (74%) | 2/6 (33%) |
| circRNA targeting reagents that do not discriminate circular and linear isoforms from the claimed host gene | 11/17 (65%) | 2/2 (100%) |
| circRNA targeting reagents **predicted to be non-targeting** in human | 6/23 (26%) | 4/6 (67%) |
| circRNA targeting reagents that could not be **verified** | 27/27 (100%) | 8/8 (100%) |
| BSJ sequence for claimed circRNA was not provided | 17/27 (63%) | 6/8 (75%) |
| Claimed circRNA sequence could not be identified in external databases | 10/27 (37%) | 2/8 (25%) |

**Table 3.** Proportions of problematic *Molecular Cancer* and *Oncogene* papers according to country of origin and institutional affiliation type

|  | *Molecular Cancer* |  |  | *Oncogene* |  |  |
| --- | --- | --- | --- | --- | --- | --- |
| **Country of origin** | **All problematic papers** | **Hospital affiliated** | **Not hospital affiliated** | **All problematic papers** | **Hospital affiliated** | **Not hospital affiliated** |
| **All countries** | 92 (100%) | 64/92 (70%) | 28/92 (30%) | 21 (100%) | 12/21 (57%) | 9/21 (43%) |
| **China** | 68/92 (74%) | 58/68 (85%) | 10/68 (15%) | 17/21 (81%) | 12/17 (70%) | 5/17 (30%) |
| **All other countries** | 24/92 (26%) | 6/24 (25%) | 18/24 (75%) | 4/21 (19%) | 0/4 (0%) | 4/4 (100%) |
| *USA* | 7/92 (8%) | 0/7 | 7/7 | - | - | - |
| *Germany* | 4/92 (2%) | 2/4 | 2/4 | - | - | - |
| *France* | 3/92 (3%) | 1/3 | 2/3 | - | - | - |
| *India* | 2/92 (2%) | 0/2 | 2/2 | - | - | - |
| *Spain* | 2/92 (2%) | 1/2 | 1/2 | - | - | - |
| *Italy* | 1/92 (1%) | 0/1 | 1/1 | - | - | - |
| *Japan* | 1/92 (1%) | 0/1 | 1/1 | 1/21 (5%) | 0/1 | 1/1 |
| *Singapore* | 1/92 (1%) | 0/1 | 1/1 | - | - | - |
| *Taiwan* | 1/92 (1%) | 0/1 | 1/1 | 1/21 (5%) | 0/1 | 1/1 |
| *Thailand* | 1/92 (1%) | 0/1 | 1/1 | - | - | - |
| *United Kingdom* | 1/92 (1%) | 0/1 | 1/1 | 1/21 (5%) | 0/1 | 1/1 |
| *Belgium* | - | - | - | 1/21 (5%) | 0/1 | 1/1 |

**Table 4.** Post-publication notices and PubPeer commentary for Molecular Cancer and Oncogene papers *Some papers were corrected/ retracted/ flagged on PubPeer due to multiple issues. **Papers corrected or flagged on PubPeer because of wrongly identified sequences include 3 previously reported *Molecular Cancer* papers [19, 21].

|  | *Molecular Cancer* | *Oncogene* |
| --- | --- | --- |
| **Problematic papers with wrongly identified nucleotide sequence (s)**<br>**(proportion (percentage))** | **92/92 (100%)** | **21/21 (100%)** |
| **Published corrections\*** | **11/92 (12%)** | **4/21 (19%)** |
| Images | 10/11 (91%) | 3/4 (75%) |
| Wrongly identified nucleotide sequences** | 2/11 (18%) | - |
| Typographical errors | 1/11 (9%) | - |
| Statistics | - | 1/4 (25%) |
| **Retractions\*** | **4/92 (4%)** | - |
| Images | 4/4 (100%) | - |
| Ethics | 2/4 (50%) | - |
| **Flagged on PubPeer\*** | **27/92 (29%)** | **5/21 (24%)** |
| Images | 24/27 (89%) | 4/5 (80%) |
| Wrongly identified nucleotide sequences** | 4/27 (15%) | - |
| Methodology | 2/27 (7%) | 2/5 (40%) |
| Ethics | 1/27 (4%) | - |
| Concerns about published correction | 1/27 (4%) | - |
| Duplicate publication | 1/27 (4%) | - |
| **Papers describing non-verifiable circRNA targeting reagent(s)** | **5/5 (100%)** | - |
| **Published corrections** | **1/5 (20%)** | - |
| Typographical error (circRNA identifier) | 1/1 (100%) | - |

### Targeted *Oncogene* corpus

To investigate whether original papers with wrongly identified or non-verifiable nucleotide sequences can be identified in other high IF cancer research journals, we verified nucleotide sequence reagent identities in a subset of original *Oncogene* papers. As described in the Methods, we employed keyword-driven searches of *Oncogene* papers published in 2020, using keywords identified in some problematic *Molecular Cancer* papers (miRNA, miR, circular RNA, or circRNA). This search strategy identified a corpus of 52 *Oncogene* papers that commonly described the analysis of one or more miR’s and/or circRNAs (Table 1). Most (42/52, 81%) selected *Oncogene* papers described human research and at least one nucleotide sequence that was claimed to target a non-modified human gene or genomic sequence. These 42 papers described a median number of 20 sequences/ paper (range 2-115) (Table 1).

### Problematic *Oncogene* papers describing wrongly identified nucleotide sequence(s)

The 42 *Oncogene* papers included 1,165 nucleotide sequences, of which 50 (4.3%) sequences were predicted to be wrongly identified (Table 1, Table S1). These 50 wrongly identified sequences were distributed across 21/52 (40%) corpus papers and 21/42 (50%) screened papers (Table S2). The 21 problematic *Oncogene* papers described a median of 2 wrongly identified sequences/ paper (range 1-5) (Table 1). Problematic *Oncogene* papers described experiments in human cancer models that corresponded to 14 different cancer types, most frequently breast cancer and hepatocellular carcinoma (Table S2). Problematic *Oncogene* papers referred to a median of 3 genes or transcripts in their titles (range 0-4), where most titles referred to miR(s) (13/21, 62%) (Table S2).

As in problematic *Molecular Cancer* papers, most wrongly identified sequences in 2020 *Oncogene* papers represented targeting reagents that were verified to target a different human gene or genomic sequence from that claimed (27/50, 54%), followed by claimed targeting reagents that were predicted to be non-targeting in human (23/50, 46%) (Table 1). Six wrongly identified sequences were claimed to target human circRNAs, which were either predicted to be non-targeting in human or to target linear transcript(s) from the claimed host gene (Table 2). A further 8 circRNA-targeting sequences were not verifiable, either because the relevant BSJ sequence was not provided or the claimed circRNA sequence could not be identified (Table 2, Tables S3, S5).

### Countries of origin and institutional affiliations of problematic *Molecular Cancer* and ***Oncogene* papers**

Problematic *Molecular Cancer* and *Oncogene* papers were authored by teams from 12 and 5 different countries, respectively (Table 3, Table S2). Most problematic *Molecular Cancer* (68/92, 764%) and *Oncogene* papers (17/21, 81%) were authored by teams from China, followed by authors from USA in the case of *Molecular Cancer* papers (7/92, 8%) (Table 3). When problematic papers were analysed according to both country and institution of origin [21], most problematic *Molecular Cancer* and *Oncogene* papers from China were affiliated with hospitals, compared with minorities of problematic papers from other countries (Table 3). Significantly more problematic *Molecular Cancer* papers from China were authored by hospital-affiliated teams (58/68 (85%)), compared with problematic papers from other countries (6/24 (25%)) (Fisher’s exact test, p<0.0001, n=92) (Table 3).

### Citations and post-publication commentary/ corrections of problematic *Molecular Cancer* and *Oncogene* papers

The 92 problematic *Molecular Cancer* papers have been collectively cited 8,048 times according to Google Scholar, including PubMed ID 32384893, which has been cited 240 times since publication in 2018. Some 34 problematic *Molecular Cancer* papers have been cited at least 100 times, and 27 others have been cited at least 50 times (Fig. 7). Highly cited papers include 22 papers published in 2020 (Fig. 7). The 21 problematic 2020 *Oncogene* papers have been cited 878 times according to Google Scholar, where one paper has been cited 168 times, and 5 other papers have been cited at least 50 times (Fig. 7).

**Figure 7.**
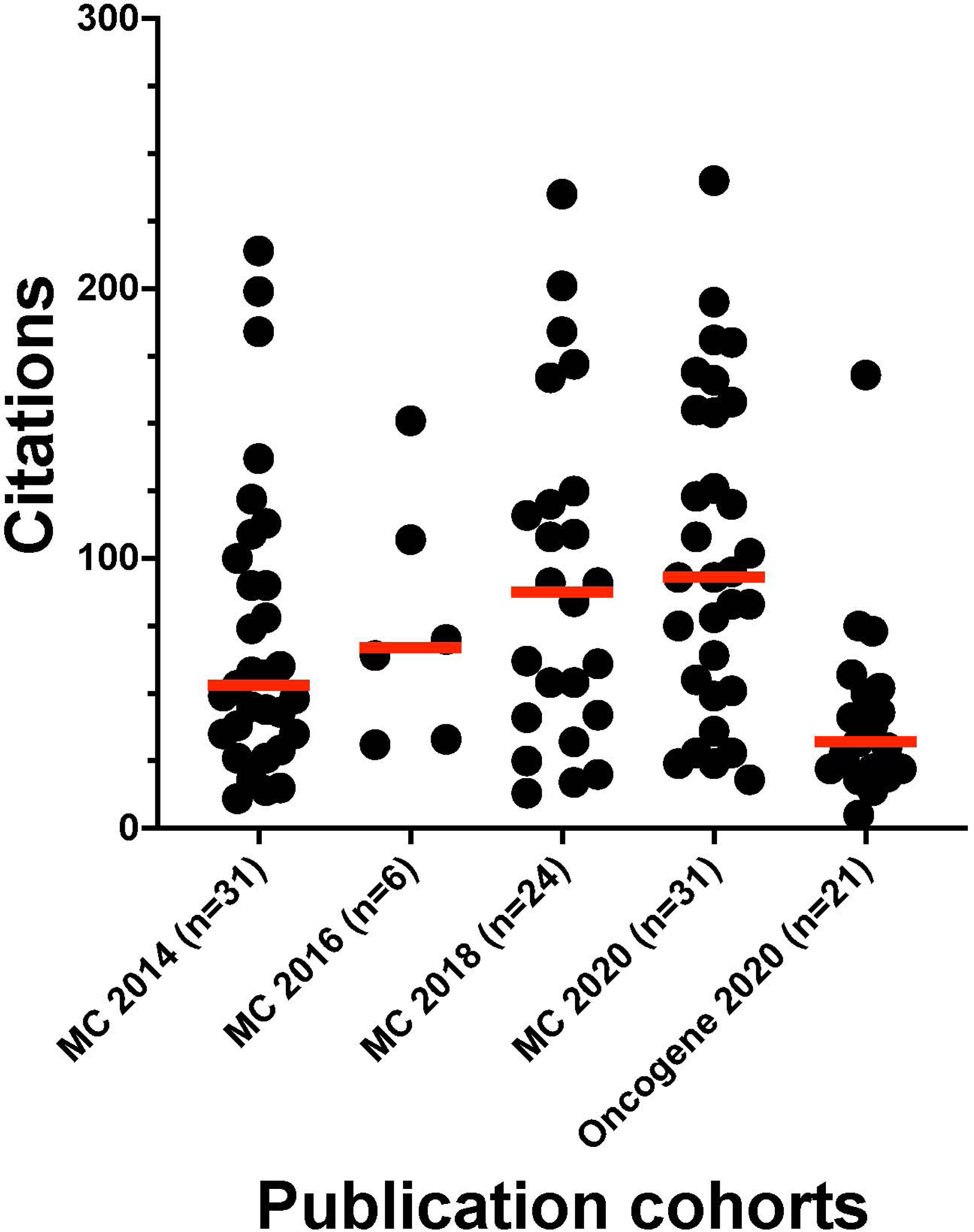
Google Scholar citations of problematic *Molecular Cancer* and *Oncogene* papers (Y axis) according to journal and publication year (X axis). Individual/ median citation numbers are shown as black dots/ red horizontal lines, respectively. Numbers of problematic *Molecular Cancer* (MC) or *Oncogene* papers per year are shown below the X-axis.

Eleven problematic *Molecular Cancer* papers, 4 problematic *Oncogene* papers, and one *Molecular Cancer* paper with non-verifiable circRNA targeting reagents have associated published corrections, mostly in response to concerns about image integrity (Table 4). Two *Molecular Cancer* papers were corrected for wrongly identified sequences (Table S6), where one paper had been previously identified by our team [21]. In the other published correction, one nucleotide sequence remained wrongly identified (Table S6). Four problematic *Molecular Cancer* papers have been retracted in response to image integrity and ethics concerns (Table 4). Just under one third (27/92, 29%) of problematic *Molecular Cancer* papers and 5/21 (24%) *Oncogene* papers have been flagged on PubPeer [43], mostly for image integrity concerns (Table 4). Four problematic *Molecular Cancer* papers have been flagged on PubPeer for wrongly identified nucleotide sequences, including one paper from a previous study [19] (Table 4).

## Discussion

Verifying the identities of nucleotide sequences published in Molecular Cancer has shown that 10-38% of all original *Molecular Cancer* papers published in 2014, 2016, 2018 and 2020 papers described wrongly identified nucleotide sequence(s). The proportions of problematic *Molecular Cancer* papers also rose from 2014-2020, when the journal IF increased from 4.3-27.3 (Figure 1). We identified similar problematic papers in the journal *Oncogene*, where 40% of 2020 *Oncogene* papers that studied miRs and/or circRNAs were found to describe wrongly identified nucleotide sequence(s). Many problematic papers in both *Molecular Cancer* and *Oncogene* have been highly cited, including recent publications from 2020. These results support and extend previous findings demonstrating that human gene research papers with wrongly identified nucleotide sequences can be identified in high IF journals [19, 21].

The analysis of *Molecular Cancer* and *Oncogene* papers that examined circRNAs in human cancer also identified incorrect circRNA targeting reagents, where some errors reflected the particular requirements of circRNA-targeting reagents [44–46]. As also reported by Zhong et al. [47], we identified claimed divergent RT-PCR primers that did not appear to discriminate between circular and linear transcripts, as well as single reagents that did not appear to be specific for the claimed circRNA target. The identities of other circRNA-targeting reagents could not be verified, either because the claimed circRNA sequence or the BSJ sequence was not provided and/or could not be identified elsewhere. These results add to previous descriptions of cancer research papers in which the identities of claimed circRNAs could not be independently verified [48].

Before discussing our results further, it is important to recognize our study’s limitations, as well as study design factors that may have identified higher numbers of problematic papers than those previously reported [21]. We recognize that the present study has only examined original papers from two journals, due to the challenges of manually verifying nucleotide sequence identities in papers that frequently described 50-100 sequences per paper. In previous studies, we employed the semi-automated Seek & Blastn tool [19], which screens publications for short nucleotide sequences and then verifies their claimed identities using blastn [49]. Screening original papers with Seek & Blastn and then manually verifying the results found that up to 4.2% and 12.6% of 2014-2018 papers in the journals *Gene* and *Oncology Reports* described wrongly identified nucleotide sequence(s). In the present study, every *Molecular Cancer* and *Oncogene* paper was analyzed manually, which may have reduced false-negative results associated with Seek & Blastn screening [19, 21].

The numbers of nucleotide sequences per *Molecular Cancer* paper also rose significantly from 2014 to 2020, where we noted a positive correlation between the numbers of wrongly identified sequences and the numbers of sequences analyzed per paper. It is therefore possible that as the numbers of nucleotide sequence reagents per paper increase, more papers could describe small numbers of wrongly identified sequences. At the same time, the median numbers of wrongly identified sequences per problematic *Molecular Cancer* paper were largely stable across 2014-2020. Median numbers of wrongly identified sequences in problematic *Molecular Cancer* and *Oncogene* papers were also similar to those noted for problematic papers in lower IF journals [21]. This suggests that the rising proportions of problematic *Molecular Cancer* papers from 2014-2020 do not simply reflect the publication of increasingly complex papers during this time.

Clearly, wrongly identified nucleotide sequences can occur in the context of genuine research [21], particularly where papers describe many individual reagents. At the same time, many of the nucleotide sequence identity errors in *Molecular Cancer* and *Oncogene* papers seem implausible. These include claimed human gene targeting sequences with no identifiable human target, where some sequences were instead predicted to target orthologous genes in species other than human. As we have previously described, most researchers will be unlikely to select human gene targeting reagents that do not target any human gene [21]. Such reagents would be typically flagged prior to publication by generating uniformly negative results. Similarly, most researchers will be aware that nucleotide sequence reagents that are identical to gene sequences in rodents, plants or fungi will be very unlikely to effectively target the orthologous human gene [21]. It is also notable that some problematic *Molecular Cancer* and *Oncogene* papers have either been corrected, retracted and/or flagged on PubPeer, most frequently for image integrity issues.

These shared and implausible features of many problematic *Molecular Cancer* and *Oncogene* papers could add to journal concerns [34–36] that paper mills may be successfully targeting some high IF journals. Given the prestige associated with publishing in high IF journals, it seems likely that some paper mills and their clients will value or require publications in high IF journals. This requirement may become acute as lower IF journals are recognized as possible paper mill targets [50]. As the price per paper mill manuscript may be partly dictated by journal IF [51], publishing in high IF journals could allow paper mills to charge higher manuscript fees, which could allow paper mills to produce more sophisticated manuscripts that more closely resemble genuine papers. Developments in artificial intelligence, both in terms of text [52, 53] and image generation [54, 55], could add to paper mill capacity to produce sophisticated manuscripts that could meet the expectations of some high IF journals.

Researchers read the scientific and scholarly literature for many purposes, including education, general interest, and to find new ideas and topics for research [56]. Due to limitations in available time and human cognition, academics and researchers have consistently described reading between ∼150-400 research publications per year [56–58]. As human reading capacity is greatly exceeded by the quantity of available literature, many researchers use heuristics to help decide which papers they should read [59–62]. Survey results consistently report that academics and researchers prioritize reading papers in high IF journals and/or with high citation numbers [59, 60, 62]. Furthermore, younger researchers may place more emphasis on journal IF and citations as proxies for research quality [59, 60].

The repeated demonstration of researcher preferences for papers in high IF journals [59, 60, 62] means that significant proportions of problematic papers in high IF cancer journals could seriously impact future research. Highly cited papers in high IF journals are likely to be prioritized for reading [59, 60, 62], where a proportion of these papers will be used in future research. Researchers may also be more motivated to reproduce results published in high IF journals, as reflected by the design of the Cancer Biology Reproducibility Project that attempted to reproduce cancer research studies published in high IF journals [4]. Problematic gene research papers in high IF cancer journals could therefore encourage more researchers to attempt new research based on problematic results, and to waste time and resources through the experimental use of wrongly-identified reagents [15, 21].

Due to the direct relationship between citations and journal IF, citations to problematic papers could also be generating a positive feed-back loop within the human gene literature. Highly cited problematic papers could boost journal IF, which could also bring these papers to the attention of more researchers who use journal IF and citation numbers as proxies for research quality [59, 60]. Editor awareness that papers about ncRNAs attract high numbers of citations [63] could also lead journal editors to prioritize these manuscripts, as highly cited papers will boost their journal’s IF. The unfortunate confluence between high citations of ncRNA publications [63] and the possible value of these gene topics to paper mills [15, 17, 21, 31, 39] could drive both the acceptance of problematic human gene research manuscripts by high IF journals, and then bring these publications to the attention of more researchers.

The identification of superficially plausible yet problematic papers in high IF cancer research journals should encourage the analysis of recent papers in other high IF journals. Problematic papers in high IF journals could demonstrate the leading edge of paper mill capability and help to predict the types of manuscripts that could be received by a broader range of journals in future [15]. The possibility of paper mills harnessing new and rapidly developing capacities for automated text generation [53] highlights the urgent need for more critical analyses of papers in high IF journals.

The field of circRNA research is also growing rapidly, where the overwhelming majority of circRNA papers have been published by authors from few countries [64, 65]. Our results indicate that circRNAs may provide a new category of human gene topics that could be exploited by paper mills. Incomplete and non-overlapping circRNA databases [66, 67] that can include poorly or incompletely annotated circRNA sequences [66, 68], combined with multiple circRNA nomenclature systems [46, 66–68] can collectively underpin superficial published descriptions of individual circRNAs, and render poor quality circRNA research more challenging to detect. Individual circRNAs can also be linked many different protein-coding genes and ncRNAs [68, 69], which could enable the creation of large numbers of manuscripts that combine different circRNAs, ncRNAs and other genes or proteins across different diseases such as human cancer types. The rapid growth in the numbers of circRNA papers [64, 65, 68] could also limit the availability of expert peer reviewers with in-depth knowledge of critical factors in circRNA research.

Regardless of the origins of the problematic circRNA papers that we have identified, our analyses show that some human circRNA papers in high IF journals are setting poor standards for methods and results reporting, particularly for readers who may be unfamiliar with the requirements of circRNA-targeting reagents. Some descriptions of circRNA research in *Molecular Cancer* and *Oncogene* indicate the need for better reporting of circRNAs and their targeting reagents (Table 5), as also recognized by others [46, 48, 66–69]. The poor reporting practices that we and others have identified (Table 5) indicate the need for guidance around circRNA (reagent) reporting to be more strictly enforced by journals, where high IF journals are well placed to show leadership on best practices.

**Table 5.** Recommendations for improved reporting of circRNA sequences and targeting reagents in research publications

| Problems | Proposed solutions |
| --- | --- |
| • Non-verifiable circRNA targeting reagents as claimed circRNA and/or sequence could not be identified [48]<br>• Identities of claimed circRNA targets unclear or poorly described [66]<br>• Case sensitive circRNA identifiers | • circRNA sequences to be described in publications and deposited in external databases with clearly disclosed accession information [66, 68]<br>• circRNA sequence descriptions to specify whether sequence is complete or partial<br>• circRNAs to be identified by unique, recognised identifiers that disclose the host gene [66, 67, 69]<br>• circRNA identifiers to be accompanied by circRNA genomic co-ordinates, including reference genome build [46, 67]<br>• Link circRNA identifiers to database entries in publications<br>• Build circRNA database search algorithms that accept case-insensitive circRNA identifiers as queries |
| • circRNA reagents could not be verified as BSJ sequence not provided or traceable [66]<br>• Limited BSJ sequence information provided in publications<br>• BSJ sequences shown in images, ie flat files, not machine readable | • circRNA descriptions to include transparent and verifiable information about BSJ sequence [66, 68]<br>• circRNA databases to annotate BSJ at sequence level and define whether BSJ is predicted or experimentally verified<br>• Require informative BSJ sequences within publications, eg. 10 nts on either side of BSJ<br>• Require BSJ sequences to be specified in machine-readable format, preferably within the main text of publication |
| • Unclear targeting parameters for single circRNA targeting reagents<br>• circRNA targeting RT-PCR primers not specified as divergent or convergent | • Single circRNA reagent descriptions to specify whether reagent targets the BSJ, or other circRNA feature that is not conserved in linear transcripts<br>• circRNA reagent descriptions to specify intended experimental use, including reagents described in supplementary tables/ files |

In summary, despite well-recognized limitations in the use of journal IF to predict research quality [41, 70], high IF journals are valued and relied upon by many biomedical researchers. Our results indicate that contrary to reasonable expectations, problematic gene research papers may be frequent in some high IF cancer journals. This highlights the need for biomedical researchers to exercise extreme caution when interpreting published gene research, including research published in high IF journals. Publications must not be exempt from critical analysis simply because they have been published in a high IF journal and/or achieved seemingly impressive numbers of citations.

Misplaced beliefs that paper mills are only a problem for lower IF journals risk exacerbating the vulnerability of high IF journals towards paper mills. Given their established brands, reputations, and available resources, we hope that high IF journals and their publishers will be responsive to reports of problematic gene research papers and will lead efforts in recognizing and responding to the threats posed by research paper mills.

## Methods

### Identification of literature corpora

*Molecular Cancer* papers were retrieved via the Web of Science using the search criteria: PY = “2014, 2016, 2018, 2020”, SO = “MOLECULAR CANCER”, AND DT = “Article”. Article titles were used as search queries on the *Molecular Cancer* website to obtain pdfs and supplementary files. Based on features of some problematic *Molecular Cancer* papers, selected *Oncogene* papers were retrieved via the Web of Science using the search criteria: PY = “2020”, SO = “ONCOGENE”, DT = “Article” and keywords = [(“Circular RNA*.mp.” OR “circRNA*.mp.”) OR (“microRNA*.mp. OR “miR*.mp.”)]. *Oncogene* article titles were used as search queries to obtain article pdfs and supplementary files through the University of Sydney library.

### Visual inspection and screening

Each article was subjected to visual screening and considered eligible for analysis if the study described the sequence of at least one nucleotide sequence reagent that was claimed to target an unmodified human transcript or genomic region. Papers including supplementary files were visually inspected to determine the claimed genetic and/or experimental identity of each nucleotide sequence. If the claimed target or experimental use of any sequence was not evident, or if a sequence was claimed to target a species other than human, the sequence was excluded from further analysis. We included papers with post-publication notices such as retractions and published corrections, except where post-publication corrections had corrected all wrongly identified nucleotide sequences at the time of publication screening.

Eligible papers were identified by their PMIDs, and nucleotide sequences and their claimed identities were extracted from text and/or supplementary files and recorded in Microsoft Excel.

### Manual verification of nucleotide sequence reagent identities

Nucleotide sequence reagents that were claimed to target human protein-coding genes and microRNAs were analysed as described [21, 71]. GeneCards [72] and GenBank [73] were used to clarify synonymous human gene identifiers. For nucleotide sequence reagents that were claimed to target long non-coding RNAs (lncRNAs), the claimed identifier was searched on lncBASE [74] and GeneCards [72] to identify the genomic co-ordinates of the claimed lncRNA. Claimed targeting reagent sequences were queried using BLAT [75] against the GRCh38/hg38 assembly and using blastn as described [21].

Nucleotide sequence reagents that were claimed to target genomic sequences including gene promoters were queried using BLAT [75] against the GRCh38/hg38 assembly as described [21]. Claimed gene promoter targeting reagents were accepted as targeting if these reagents mapped within 100 kb upstream of the claimed target gene and if reagents did not include coding gene exons. Where the claimed reagent identity did not match the verified identity, sequences were queried using BLAT [75] against earlier human genome assemblies.

### Manual verification of claimed circular RNA (circRNA) targeting reagents Verification of RT-PCR primers claimed to target circRNAs

circRNAs are alternatively-spliced transcripts where gene exons are joined through back-splicing to create circular transcripts [44–46]. RT-PCR amplification of circRNAs requires two sets of RT-PCR primers. Divergent RT-PCR primers are used to amplify the claimed circRNA by facing towards and amplifying across the circRNA BSJ [44–47]. Divergent RT-PCR primers should therefore not amplify linear transcripts from the host or any other human gene. In contrast, convergent RT-PCR primers are employed to amplify linear transcripts, typically from the claimed host gene [44–47].

For claimed divergent RT-PCR primers, forward and reverse primers were first queried on circPRIMER [45] using standard settings (Fig. 3). RT-PCR primers were accepted as correctly targeting if circPRIMER aligned both RT-PCR primer sequences to the claimed circRNA(s), such that RT-PCR primers faced towards and were predicted to amplify the BSJ (Fig. 3). If circPRIMER analyses produced no output, we then checked whether the claimed circRNA was indexed by a publicly available circRNA database such as circBASE [76] or circATLAS [77] through the disclosure of a specific circRNA identifier, or if the circRNA sequence and/or its genomic sequence co-ordinates were disclosed by the authors. If the claimed circRNA could not be identified, the claimed divergent RT-PCR primers were classified as non-verifiable [48]. If the claimed circRNA could be identified but the BSJ could not be predicted, claimed divergent RT-PCR reagents were also classified as non-verifiable.

If the claimed BSJ sequence was either disclosed or the associated genomic co-ordinates could be predicted, divergent RT-PCR primers were then queried either using the BLAT function of circBASE [76], manually mapped to the claimed circRNA sequence, and/or queried using BLAT against the GRCh38/hg38 genomic assembly. Claimed divergent RT-PCR primers were classified as wrongly identified if they did not amplify the (predicted) BSJ (Fig. 3). Wrongly identified RT-PCR primers were subjected to further analyses to classify these reagents according to nucleotide sequence error categories (see below), as described [21]. Claimed convergent RT-PCR primers were verified as previously described for RT-PCR primers targeting linear transcripts [19–21, 71].

### Verification of single nucleotide sequence reagents claimed to target circRNAs

Single reagents such as si/shRNAs and other oligonucleotides acquire circRNA specificity by targeting specific BSJ sequences [44, 46]. We first determined whether the claimed circRNA was indexed in a publicly available circRNA database, as described above, and whether the BSJ sequence could be identified (Fig. 4). If claimed circRNA or the BSJ sequence could not be identified, reagents were classified as non-verifiable (Fig. 4).

Verifiable single reagents were then aligned against the claimed circRNA sequence including the BSJ sequence (Fig. 4). Single reagents were classified as correctly targeting if they showed 100% identity to the claimed BSJ sequence and 5-16 nucleotides of sequence identity to circRNA sequences on either/ both sides of the BSJ [44]. If a claimed circRNA targeting reagent showed 100% identity to 17 or more consecutive nucleotides within any host gene exon, the reagent was classified as wrongly identified, as such reagents would not be predicted to discriminate between circular and linear transcripts from the host gene. All wrongly identified single circRNA-targeting reagents were then subjected to further verification analyses as described [21, 71].

### Classification of wrongly identified reagents according to error categories

Wrongly identified nucleotide sequence reagents were classified according to previously described error categories, namely (i) claimed targeting reagents that were predicted to target another human gene or genomic sequence, (ii) claimed targeting reagents that were predicted to be non-targeting in human and (iii) claimed non-targeting reagents that were predicted to target a human gene or transcript [19–21]. Claimed circRNA-targeting reagents (divergent RT-PCR primers, si/shRNAs, molecular probes) that were predicted to (also) target linear transcripts (from the host gene) were classified as targeting a different gene/ transcript from that claimed (category (i) above).

### Additional publication analyses

For each eligible article, we recorded the number and proportion of wrongly identified nucleotide sequence reagents. We also recorded the numbers and identities of non-verifiable circRNA reagents, noting that we did not categorise non-verifiable reagents as wrongly identified. Publications were described as problematic if they included at least one wrongly identified nucleotide sequence reagent. Papers that described non-verifiable circRNA targeting reagent(s) but no wrongly identified nucleotide sequences were reported separately. Proportions of problematic papers/ papers analysed, problematic papers/ papers screened and wrongly identified nucleotide sequences/ nucleotide sequences analysed were calculated for journals and publication years using MS Excel.

Problematic publication titles were visually inspected to identify human gene or transcript identifiers, human cancer types, and drug identifiers which were confirmed through Google searches. Human genes were categorized as either protein-coding or ncRNAs according to GeneCards [72]. The country of origin and institutional affiliation of each problematic article were identified as described [21]. Where there was no numeric majority, the first author’s affiliation was used to decide the country of origin and/or institutional affiliation. PubPeer notifications of problematic papers were identified on 16 January 2023. Reported numbers of post-publication notices linked with problematic papers are those identified through PubMed and Google Scholar searches conducted on 17 January 2023. Citations of problematic papers according to Google Scholar were collected on 22 January, 2022.

### Statistics analyses

Fisher’s exact tests conducted on GraphPad PRISM compared proportions of problematic *Molecular Cancer* papers according to publication year, and countries and institutions of origin. Shapiro-Wilk’s test was used to test for normality, and the Mann-Whitney test was conducted to compare median numbers of wrongly identified sequences per *Molecular Cancer* article according to publication year. For all problematic papers in *Molecular Cancer*, Spearman’s rank correlation coefficient was calculated between the numbers of wrongly identified sequences and numbers of analysed nucleotide sequences per article. Graphs were produced on GraphPad PRISM 9.2.

## Acknowledgements

JAB gratefully acknowledges funding from the National Health and Medical Research Council of Australia (NHMRC) Ideas grant ID APP1184263, and from the Faculty of Medicine and Health at the University of Sydney. PP is supported by a Research Training Program scholarship at the University of Sydney. We thank Dr Thomas Stoeger and Mr Reese Richardson (Northwestern University, USA) for critical reading and discussions, and Prof Lenka Munoz (University of Sydney, Australia), Prof Cyril Labbé (Univ. Grenoble Alpes, France) and Prof Guillaume Cabanac (Univ. Toulouse, France) for discussions. We apologize to authors whose work was not cited due to limits in the number of references that could be cited.

## Author contributions

Conceptualization: JAB; Methodology: PP, FJE, YP, JAB; Formal analysis: PP, YP, JAB; Writing - original draft preparation: PP, JAB; Writing - review and editing: JAB, PP, FJE, YP; Funding acquisition: JAB, PP; Supervision: JAB

## Data availability statement

All data are described in the manuscript and supplementary files. Google Scholar citation data, and PubPeer notifications are available within the public domain.

## Additional Information (including a Competing Interests Statement)

The authors have no relevant financial or non-financial interests to disclose.

## References

1. Bowen, A. & Casadevall, A. Increasing disparities between resource inputs and outcomes, as measured by certain health deliverables, in biomedical research. Proc. Natl. Acad. Sci. USA. 112, 11335–11340 (2015).

2. Pusztai, L., Hatzis, C. & Andre, F. Reproducibility of research and preclinical validation: problems and solutions. Nat. Rev. Clin. Oncol. 10, 720–724 (2013).

3. Mobley, A., Linder, S. K., Braeuer, R., Ellis, L. M. & Zwelling, L. A survey on data reproducibility in cancer research provides insights into our limited ability to translate findings from the laboratory to the clinic. PloS ONE 8, e63221 (2013).

4. Errington, T. M., Denis, A., Perfito, N., Iorns, E. & Nosek, B. A. Challenges for assessing replicability in preclinical cancer biology. Elife 10, e67995 (2021).

5. Smaldino, P. E. & McElreath, R. The natural selection of bad science. R. Soc. Open Sci. 3, 160384 (2016).

6. Kaelin, W. G. Jr. Common pitfalls in preclinical cancer target validation. *Nature Rev*. Cancer 17, 425–440 (2017).

7. Brown, A. W., Kaiser, K. A. & Allison, D. B. Issues with data and analyses: Errors, underlying themes, and potential solutions. Proc. Natl. Acad. Sci. USA 115, 2563–2570 (2018).

8. Stroebe, W., Postmes, T., & Spears, R. Scientific misconduct and the myth of self-correction in science. Perspect. Psychol. Sci. 7, 670–688 (2012).

9. Gopalakrishna, G., Ter Riet, G., Vink, G., Stoop, I., Wicherts, J.M. & Bouter L.M. Prevalence of questionable research practices, research misconduct and their potential explanatory factors: A survey among academic researchers in The Netherlands. PLoS One 17, e0263023 (2022).

10. Parker, L., Boughton, S., Lawrence, R. & Bero, L. Experts identified warning signs of fraudulent research: a qualitative study to inform a screening tool. J. Clin. Epidemiol. 151, 1–17 (2022).

11. Byrne, J. A., Grima, N., Capes-Davis, A. & Labbé, C. The possibility of systematic research fraud targeting under-studied human genes: causes, consequences and potential solutions. Biomarker Insights 14, 1–12 (2019).

12. Byrne, J. We need to talk about systematic fraud. Nature. 566, 9 (2019).

13. Byrne, J. A. & Christopher, J. Digital magic, or the dark arts of the 21st century-how can journals and peer reviewers detect manuscripts and publications from paper mills? FEBS Lett. 594, 583–589 (2020).

14. COPE. & STM. Paper Mills -research report from COPE & STM -English. https://doi.org/10.24318/jtbG8IHL (2022).

15. Byrne, J. A., Park, Y., Richardson, R. A. K., Pathmendra, P., Sun, M. & Stoeger, T. Protection of the human gene research literature from contract cheating organizations known as research paper mills. Nucleic Acids Res. 50, 12058–12070 (2022).

16. Tian, M., Su, Y. & Ru, X. Perish or publish in China: pressures on young Chinese scholars to publish in internationally indexed journals. Publications 4, 9 (2016).

17. Christopher, J. The raw truth about paper mills. FEBS Lett. 595, 1751–1757 (2021).

18. Byrne, J. A. & Labbé, C. Striking similarities between publications from China describing single gene knockdown experiments in human cancer cell lines. Scientometrics 110, 1471–1493 (2017).

19. Labbé, C., Grima, N., Gautier, T., Favier, B. & Byrne, J. A. Semi-automated fact-checking of nucleotide sequence reagents in biomedical research publications: the Seek & Blastn tool. PLoS One 14, e0213266 (2019).

20. Byrne, J. A., Park, Y., West, R. A., Capes-Davis, A., Cabanac, G. & Labbé, C. The thin ret(raction) line: biomedical journal responses to reports of incorrect non-targeting nucleotide sequence reagents in human gene knockdown publications. Scientometrics 126, 3513–3534 (2021).

21. Park, Y. et al. Identification of human gene research articles with wrongly identified nucleotide sequences. Life Sci. Alliance. 5, e202101203 (2022).

22. Qi, X., Deng, H. & Guo, X. Characteristics of retractions related to faked peer reviews: an overview. Postgrad. Med. J. 93, 499–503 (2017).

23. Han, J. & Li, Z. How Metrics-Based Academic Evaluation Could Systematically Induce Academic Misconduct: A Case Study. East Asian Sci. Tech. Soc. 12, 165–179 (2018).

24. Miyakawa, T. No raw data, no science: another possible source of the reproducibility crisis. Mol. Brain 13, 24 (2020).

25. Pinna, N., Clavel, G. & Roco, M. C. The Journal of Nanoparticle Research victim of an organized rogue editor network! *J*. Nanopart. Res. 22, 376 (2020).

26. Pines, J. Image integrity and standards. Open Biol. 10, 200165 (2020).

27. Hackett, R. & Kelly, S. Publishing ethics in the era of paper mills. Biol. Open 9, bio056556 (2020).

28. Seifert, R. How Naunyn-Schmiedeberg’s Archives of Pharmacology deals with fraudulent papers from paper mills. Naunyn Schmiedeberg’s Arch. Pharmacol. 394, 431–436 (2021).

29. Fisher, L. & Cox, R. RSC Advances Editorial: retraction of falsified manuscripts. RSC Adv. 11, 4194 (2021).

30. Behl, C. Science integrity has been never more important: It’s all about trust. J. Cell. Biochem. 22, 694–695 (2021).

31. Cooper, C. D. O. & Han, W. A new chapter for a better Bioscience Reports. Biosci. Rep. 41, BSR20211016 (2021).

32. De la Rosa, M.A. Steering towards success in stormy times: *FEBS Open Bio* in 2021. FEBS Open Bio. 11, 4–9 (2021).

33. Editorial. Preventing the publication of falsified research. Toxicol. Res. 10, 961 (2021).

34. Heck, S., Bianchini, F., Souren, N. Y., Wilhelm, C., Ohl, Y. & Plass, C. Fake data, paper mills, and their authors: The International Journal of Cancer reacts to this threat to scientific integrity. Int. J. Cancer. 149, 492–493 (2021).

35. Bricker-Anthony, C. & Giangrande, P. H. On integrity. Mol. Ther. Nucleic Acids 30, 595 (2022).

36. Frederickson, R. M. & Herzog, R. W. Addressing the big business of fake science. Mol. Ther. 30, 2390 (2022).

37. Kadomatsu, K. Editorial. J. Biochem. 168, 317 (2020).

38. Clark, A. J. L. & Buckmaster, S. Fake science for sale? How Endocrine Connections is tackling paper mills. Endocr. Connect. 10, E3–E4 (2021).

39. Pérez-Neri, I., Pineda, C. & Sandoval, H. Threats to scholarly research integrity arising from paper mills: a rapid scoping review. Clin. Rheumatol. 41, 2241–2248 (2022).

40. Romanovsky, M. Distribution of scientific journals impact factor. arXiv. 1904.05320 (2019).

41. Siler, K. & Larivière, V. Who games metrics and rankings? Institutional niches and journal impact factor inflation. Research Policy. 51, S0048733322001317 (2022).

42. Kempf, E. et al. Overinterpretation and misreporting of prognostic factor studies in oncology: a systematic review. Br. J. Cancer. 119, 1288–1296 (2018).

43. Barbour, B. & Stell, B.M. PubPeer: Scientific assessment without metrics. In Gaming the metrics: Misconduct and manipulation in academic research. (ed. Biagioli, M. & Lippman, A.) 149–155 (MIT Press, 2020)

44. Dudekula, D. B., Panda, A. C., Grammatikakis, I., De, S., Abdelmohsen, K., & Gorospe, M. CircInteractome: A web tool for exploring circular RNAs and their interacting proteins and microRNAs. RNA Biology 13, 34–42 (2016).

45. Zhong, S., Wang, J., Zhang, Q., Xu, H. & Feng, J. CircPrimer: a software for annotating circRNAs and determining the specificity of circRNA primers. BMC Bioinform. 19, 292 (2018).

46. Nielsen, A. F. et al. Best practice standards for circular RNA research. Nature Methods 19, 1208–1220 (2022).

47. Zhong, S. et al. Identification of internal control genes for circular RNAs. Biotechnol. Lett. 41, 1111–1119 (2019).

48. Patop, I. L. & Kadener, S. circRNAs in Cancer. Curr. Op. Genet. Dev. 48, 121–127 (2018).

49. Altschul, S. F., Gish, W., Miller, W., Myers, E. W. & Lipman, D. J. Basic local alignment search tool. J. Mol. Biol. 215, 403–410 (1990).

50. Zhang, L., Wei, Y., Sivertsen, G., & Huang, Y. The motivations and criteria behind China’s list of questionable journals. Learned Pub. 35, 467–480 (2022).

51. Abalkina, A. Publication and collaboration anomalies in academic papers originating from a paper mill: evidence from a Russia-based paper mill. Preprint at https://arxiv.org/abs/2112.13322 (2021).

52. Floridi, L. & Chiriatti, M. GPT-3: Its nature, scope, limits, and consequences. Minds Mach. 30, 681–694 (2020).

53. Grimaldi, G. & Ehrler, B. AI et al.: Machines Are About to Change Scientific Publishing Forever. ACS Energy Lett. 8, 878–880 (2023).

54. Wang, L., Zhou, L., Yang, W. & Yu, R. Deepfakes: A new threat to image fabrication in scientific publications? Patterns 3, 100509 (2022).

55. Gu, J., Wang, X., Li, C., Zhao, J., Fu, W., Liang, G. & Qiu, J. AI-enabled image fraud in scientific publications. Patterns 3, 100511 (2022).

56. Tenopir, C., King, D.W., Christian, L. & Volentine, R. Scholarly article seeking, reading, and use: a continuing evolution from print to electronic in the sciences and social sciences. Learned Pub. 28, 93–105 (2015).

57. Tenopir, C., King, D.W., Spencer, J., & Wu, L. Variations in article seeking and reading patterns of academics: What makes a difference? Lib. Inform. Sci. Res. 31, 139–148 (2009).

58. Tenopir, C., Christian, L. & Kaufman, J. Seeking, Reading, and Use of Scholarly Articles: An International Study of Perceptions and Behavior of Researchers. Publications 7, 18 (2019).

59. Tenopir, C. et al. Trustworthiness and authority of scholarly information in a digital age: Results of an international questionnaire. J. Ass. Inf. Sci. Tech. 67, 2344–2361 (2016).

60. Nicholas, D. et al. So, are early career researchers the harbingers of change? Learned Pub. 32, 237–247 (2019).

61. Morales, E., McKiernan, E. C., Niles, M. T., Schimanski, L. & Alperin, J. P. How faculty define quality, prestige, and impact of academic journals. PloS One 16, e0257340 (2021).

62. Teplitskiy, M., Duede, E., Menietti, M. & Lakhani, K. R. How status of research papers affects the way they are read and cited. Res. Policy 51, 104484 (2022).

63. Fire, M. & Guestrin, C. Over-optimization of academic publishing metrics: observing Goodhart’s Law in action. Gigascience 8, giz053 (2019)

64. Zhang, C., Kang, Y., Kong, F., Yang, Q. & Chang, D. Hotspots and development frontiers of circRNA based on bibliometric analysis. Non-coding RNA Res. 7, 77–88 (2022).

65. Wu, R. et al. Bibliometric Analysis of Global Circular RNA Research Trends from 2007 to 2018. Cell J. 23, 238–246 (2021).

66. Vromman, M., Vandesompele, J. & Volders, P. J. Closing the circle: current state and perspectives of circular RNA databases. Brief. Bioinform. 22, 288–297 (2021).

67. Costa, M. C. & Enguita F. J. Towards a universal nomenclature standardization for circular RNAs. Non-coding RNA Investig. 4, 2 (2020).

68. Dodbele, S., Mutlu, N. & Wilusz, J. E. Best practices to ensure robust investigation of circular RNAs: pitfalls and tips. EMBO Rep. 22, e52072 (2021).

69. Kristensen, L. S., Hansen, T. B., Venø, M. T. & Kjems, J. Circular RNAs in cancer: opportunities and challenges in the field. Oncogene, 37, 555–565 (2018).

70. Ioannidis, J. P. A. & Thombs, B. D. A user’s guide to inflated and manipulated impact factors. Eur. J. Clin. Invest. 49, e13151 (2019).

71. 71. Byrne J. A., Park Y., Capes-Davis, A., Favier, B., Cabanac, G. & Labbé, C. Seek & Blastn Standard Operating Procedure V.1. https://www.protocols.io/view/seek-amp-blastn-standard-operating-procedure-bjhpkj5n (2021).

72. Stelzer, G. et al. The GeneCards Suite: From Gene Data Mining to Disease Genome Sequence Analyses. Curr. Protocols Bioinf. 54, 1.30.1–1.30.33 (2016).

73. Sayers, E. W., Cavanaugh, M., Clark, K., Ostell, J., Pruitt, K. D., & Karsch-Mizrachi, I. GenBank. Nucleic Acids Res. 47, D94–D99 (2019).

74. Karagkouni, D. et al. DIANA-LncBase v3: indexing experimentally supported miRNA targets on non-coding transcripts. Nucleic Acids Res. 48, D101–D110 (2020).

75. Lee, B. T. et al. The UCSC Genome Browser database: 2022 update. Nucleic Acids Res. 50, D1115–D1122 (2022).

76. Glažar, P., Papavasileiou, P. & Rajewsky, N. circBase: a database for circular RNAs. RNA 20, 1666–1670 (2014).

77. Wu, W., Ji, P., & Zhao, F. CircAtlas: an integrated resource of one million highly accurate circular RNAs from 1070 vertebrate transcriptomes. Genome Biol. 21, 101 (2020).

